# Parallel epigenetic modifications induced by hatchery rearing in a Pacific Salmon

**DOI:** 10.1101/148577

**Authors:** Jérémy Le Luyer, Martin Laporte, Terry D. Beacham, Karia H. Kaukinen, Ruth E. Withler, Jong S. Leong, Eric B. Rondeau, Ben F. Koop, Louis Bernatchez

## Abstract

**Highlights:** - First study to highlight parallel epigenetic modifications induced by hatchery rearing as a potential explanatory mechanism for rapid change in fitness

*Summary:* A puzzling question in conservation biology is how to maintain overall fitness of individuals bred in captive environment upon release into the wild, especially for rehabilitating declining or threatened species [1,2]. For salmonid species, a heritable change in fitness related traits and gene expression has been reported to occur in a single generation of captivity in hatchery environment [3–5]. Such rapid changes are congruent with models of inadvertent domestication selection which may lead to maladaptation in the natural environment [4]. Arguably, the underlying mechanism by which captivity may induce fitness difference between wild and captive congeners is still poorly understood. Short-term selection on complex phenotypic traits is expected to induce subtle changes in allele frequency over multiple loci [7–9]. Yet, most studies investigating the molecular basis for rapid change in fitness related traits occurring in hatchery have concentrated their effort on finding evidence for selection at the genome level by identifying loci with large effect. Numerous wild stocks of Pacific anadromous salmon and trout (genus *Oncorhynchus* and *Salmo*) have experienced fluctuating abundance over the past century, with a series of sharp declines [6–8]. With the objectives of preserving ecosystem integrity, enhancing declining populations and sustaining fisheries, conservation hatcheries have been flourishing. This is particularly true along the North American Pacific coast where billions of salmonids, all species included, are released each year. Despite substantial improvement of production management, the beneficial ecological role of hatcheries in enhancing and restoring wild stocks is still debated, mainly because of the reduced fitness and maladaptation of hatchery-fish when released in the wild [3,5,9]. Although previous studies showed that domestication selection was involved in such fitness impairment, they also observed that different environmental conditions (e.g. reduced fish density) significantly modulated the physiological acclimation to hatchery environment [4]. Environmental stimuli are especially relevant during early embryonic development, which also correspond to a sensitive methylation reprogramming window in vertebrates [10,11]. It is therefore plausible that differences in rearing environment during early development may result in epigenetic modifications that could in turn impact on fitness. However, the only epigenetic study to date pertaining to captive rearing in salmonids and performed using methylation-sensitive amplified fragments (MSAP) failed to identify significant changes in methylation profile associated with hatchery rearing [12] Here, we used a higher resolution approach to compare the genome-wide pattern of methylation in hatchery-reared juvenile (smolt) Coho Salmon with that of their wild counterparts in two geographically distant rivers in British Columbia, Canada. Using a reduced representation bisulfite sequencing (RRBS) approach covering an average per individual of about 70 million cytosines in CpG context, we identified 100 methylated regions (DMRs) that differed in parallel between hatchery and natural origin salmon in both rivers. The total variance of epigenetic variation among individuals explained by river or origin and rearing environment in a RDA model was 16% (adj.R^2^=0.16), and both variables equally explained about 8% of the variance after controlling for each other. The gene ontology analysis revealed that regions with different methylation levels between hatchery and natural origin salmon showed enrichment for ion homeostasis, synaptic and neuromuscular regulation, immune and stress response, and control of locomotion functions. We further identified 15,044 SNPs that allowed detection of significant differences between either rivers or sexes. However, no effect of rearing environment was observed, confirming that hatchery and natural origin fish of a given river belong to the same panmictic population, as expected based on the hatchery programs applied in these rivers (see Supplementary experimental procedures). Moreover, neither a standard genome-scan approach nor a polygenic statistical framework allowed detection of selective effects within a single generation between hatchery and natural origin salmon. Therefore, this is the first study to demonstrate that parallel epigenetic modifications induced by hatchery rearing during early development may represent a potential explanatory mechanism for rapid change in fitness-related traits previously reported in salmonids.

## Results

### Sampling

We collected a total of 40 Coho Salmon from two rivers in British Columbia, Canada; the Capilano and Quinsam Rivers (Table S1, Figure S1). These systems are well suited to test specifically for the effect of rearing environment on patterns of methylation, independent of the genetic background between fish born in the wild (thereafter natural origin) vs. those born in hatchery (see Supplementary experimental procedures). During their downstream migration to the sea, we collected from each river 10 juveniles (smolt stage) reared in captivity in a local hatchery (fin-clipped; hereafter “hatchery origin”) and 10 smolts born in the wild (hereafter “natural origin”). Broodstock for the hatchery fish was collected while returning to spawn in the same year in both rivers. The number of returning adults sampled and used for breeding was 758 and 894 for the Capilano River and Quinsam River populations, respectively. The broodstock included 3 years old females, as well as 2-3 years old males and could represent fish born previously either in hatchery or in the wild.

### Evidence for parallel epigenetic modifications in hatchery environment

We used a Reduced Representation Bisulfite Sequencing (RRBS) approach, with the *MspI* restriction enzyme, to document both genome-wide methylation and genetic variation. In order to avoid the possibility of falsely interpreting existing C-T DNA polymorphism as epigenetic variation, we first masked the genome (GenBank assembly accession: GCA_002021735.1) by removing all C>T polymorphism (1,896,050 SNPs; maf=0.05) found by whole-genome re-sequencing of 20 fish from British Columbia (Supplementary experimental procedures; Figure S1). We used a tiling window approach to quantify the percentage of methylation over 1000-bp regions throughout the masked genome and retained only cytosines in a CpG context for downstream analyses, as these regions represent the responsive methylation context in vertebrates (Supplementary experimental procedures). We used a db-RDA to document methylation variation among hatchery and natural origin fish from both rivers. We first produced a principal coordinate analysis (PCoA) on a Euclidean distance matrix computed using all the raw data and kept axes according to the cumulative broken-stick threshold, which correspond to six axes explaining at least 2.75% of the variance for a total of 42.2% of the variance [13]. A distance-based redundancy analysis (db-RDA) was then produced on the epigenetic variation explained by these PCoA factors (response matrix) with river of origin, rearing environment and sex as explanatory variables. The model was significant with an adjusted R^2^ of 0.16 (Figure 1). Both river of origin and rearing environment were significant whereas no significant effect was detected for sex (Figure 1). Partial db-RDAs revealed that the net variation explained by rearing environment (adj.R^2^=0.08; F=4.34; p-value<0.05) was identical to the one explained by the river of origin (adj.R^2^=0.08; F=4.66; p-value<0.01). This shared variation between hatchery origin salmon from both rivers relative to their natural origin congeners provides evidence for similar (parallel) epigenetic modifications induced by hatchery rearing.

**Figure 1:**
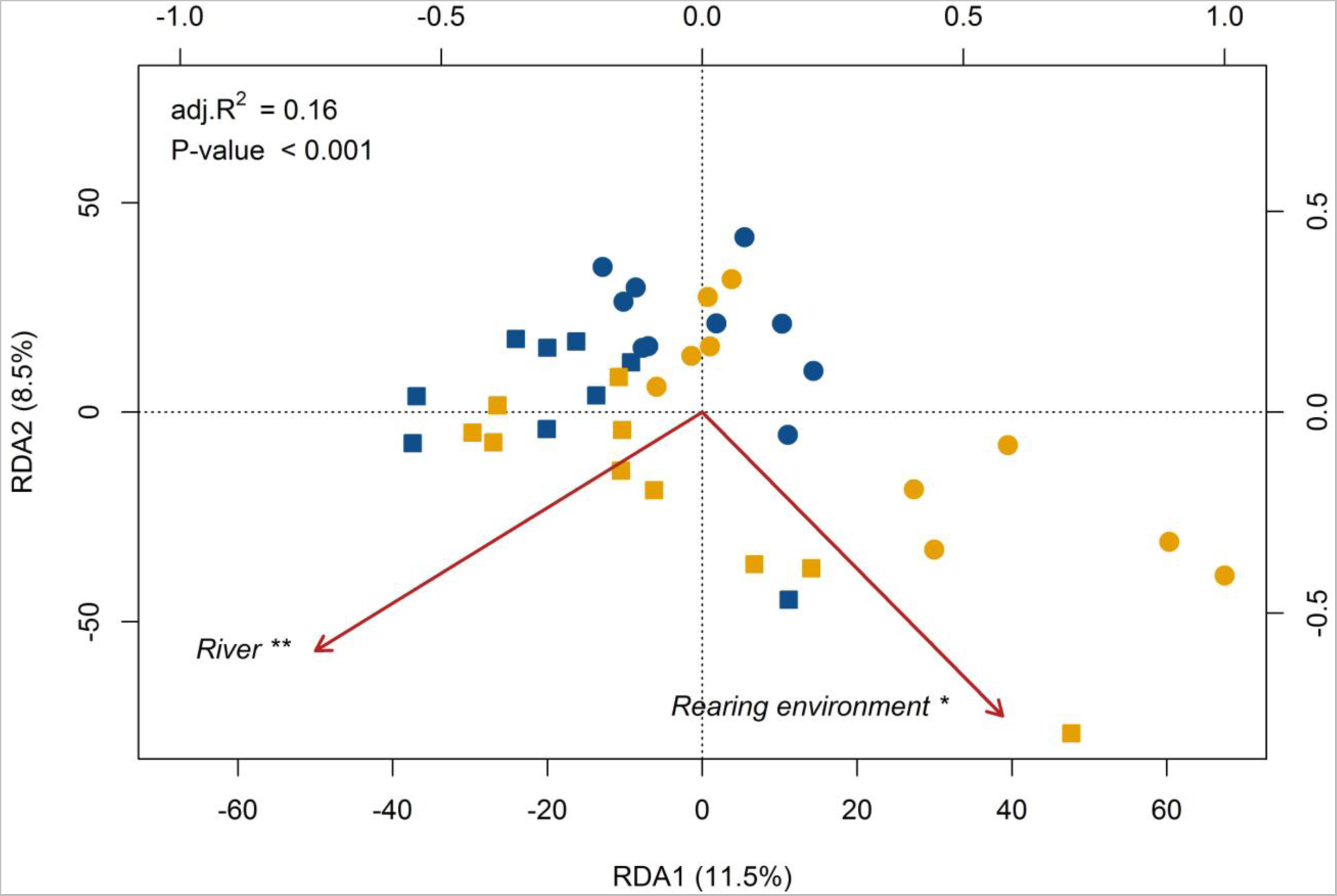
Distance-base redundancy analysis (db-RDA) performed on the DNA methylation levels of 131,807 1000-bp sliding window regions for each individual (n=39). Symbols represent rivers: Capilano (circle) and Quinsam (square). Colors represent rearing environment: hatchery (blue) and wild (yellow). The db-RDA was globally significant and explained 16% of all DNA methylation regions variation (adj.R^2^=0.16). River of origin and rearing environment both significantly explained 8% of the variation after controlling for each other with subsequent partial db-RDAs. Asterisks represent p-value < 0.01 (**) and p-value < 0.05 (*) related to the explanatory factors.

Moreover, we identified differentially methylated regions (defined as having >15% overall difference; q-value < 0.001; see Supplementary experimental procedures) between rearing environments, using a logistic regression with river of origin as covariates. We identified a total of 100 DMRs that were distributed among 27 chromosomes and 20 unmapped scaffolds (Figure 2). The proportion of hypermethylated DMRs was much greater in hatchery origin relative to natural origin salmon (89 vs 11; χ^2^=60.84, df=1, P<0.001), pointing to a general pattern of downregulation of genes associated with these DMRs in hatchery origin salmon.

**Figure 2:**
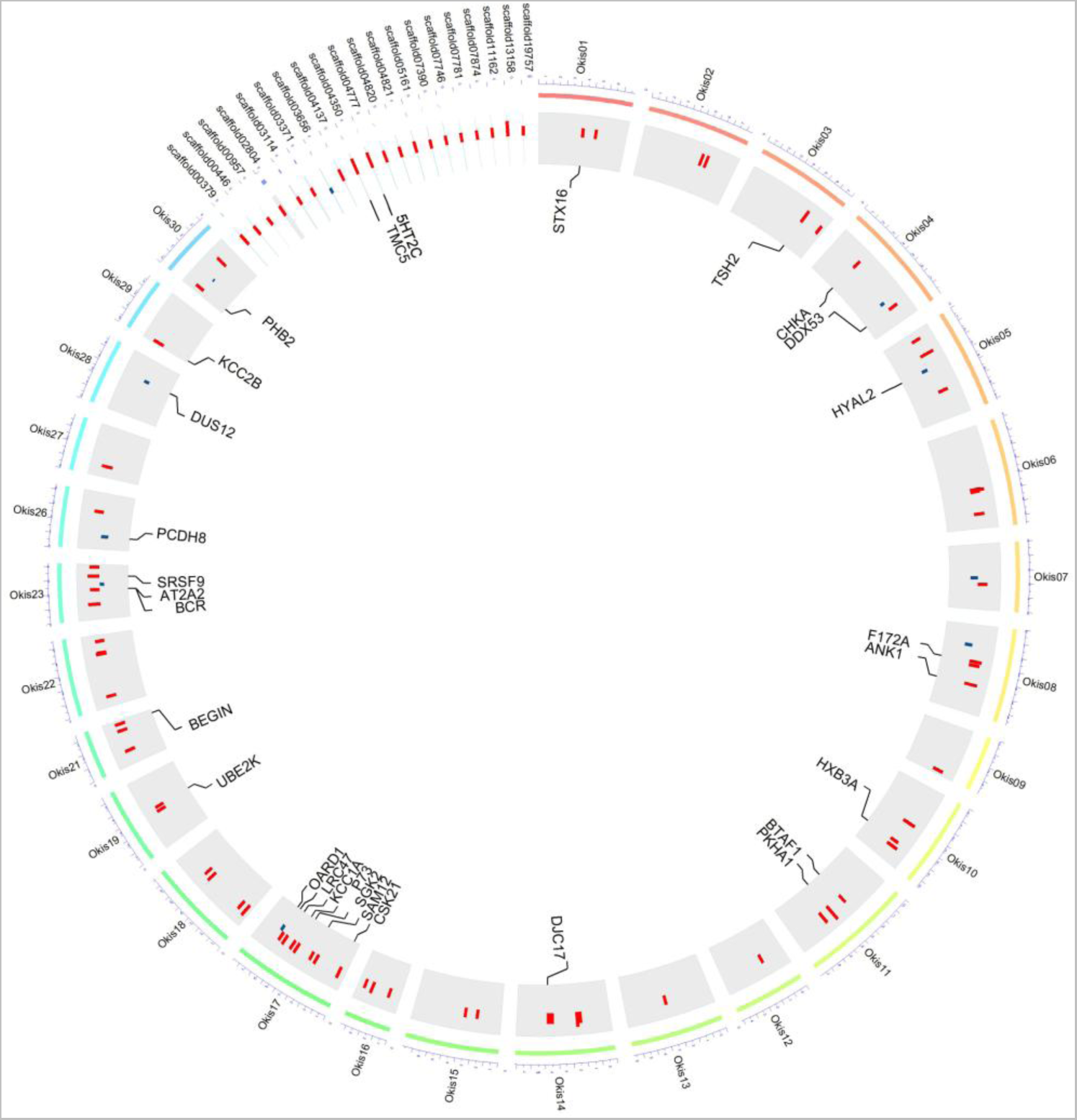
Circos plot of differentially methylated regions between hatchery and wild fish. Only the chromosomes (n = 27) and scaffolds (n = 20) containing differentially methylated regions are plotted. Barplots represent hypermethylated regions (red) and hypomethylated regions (blue) in hatchery fish. Only annotated regions (blastx e-value < 10-6) are represented.

### Functional annotation and gene ontology of DMRs

We mapped the recently published transcriptome of the Coho Salmon [14] to the species’ draft genome (see Supplemental files for details) to infer functional annotation of DMRs. Out of the 100 DMRs, we identified 37 DMRs overlapping 52 unique transcripts and regions comprising 5kb up and downstream of these transcripts. A blastx approach successfully identified 29 unique Uniprot IDs and again revealed an excess of hypermethylation in hatchery relative to wild fish (25 hypermethylated vs. 4 hypomethylated; χ^2^=15.21, df=1, P<0.001; Figure 2; Table 1). Gene ontology (GO) analysis revealed an over-representation (p-value<0.05 and at least three genes by GO term) of modules associated with ion homeostasis (GO:0055080: cation homeostasis, GO:0042592: homeostatic process, GO:0043167: ion binding, GO:0055065: metal ion homeostasis). It has been shown previously in a closely related species (Rainbow Trout, *Oncorhynchus mykiss*) that hatchery-rearing negatively affects acclimation to seawater as reflected by lower specific activity of NA+ K+ ATPase and lower survival following seawater transfer [15]. We also observed a significant enrichment for functions associated with the immune response (GO:0031347: regulation of defense response, GO:0050727: regulation of inflammatory response, GO:0045321: leukocyte activation), as well as synaptic signal modulation and locomotion functions (GO:0099572: postsynaptic specialization, GO:0050885: neuromuscular process controlling balance). The neuromuscular process controlling balance includes the calcium/calmodulin-dependent protein kinase type II subunit beta (CAMK2B), hypermethylated in hatchery fish, which is a main actor of the neuromuscular communication and regulating Ca^2+^signalling in skeletal muscle tissue [16]. Its activation has also been associated, together with the Ca^2+^signalling, to sustained and endurance muscle exercise in humans and the control of muscle development and excitation [17,18]. Lower critical swimming performance (*Uct*) has been documented in hatchery-reared Coho Salmon compared to their wild counterparts following transfer to seawater, and lower average swimming speed has been documented between wild and F1-hatchery Atlantic Salmon (*Salmo salar*) and Brown trout (*Salmo trutta*) smolts [19,20]. Moreover, the serotonin receptor 2C (HTR2C), which regulates appetite and feeding behavior, was also hypermethylated in hatchery fish [21]. Finally, we observed a GO enrichment for transcription factors (GO:0006357: regulation of transcription from RNA polymerase II promoter) which comprised the TATA-binding protein-associated factor 172, also hypermethylated in hatchery-origin fish, which is involved in the global transcription regulation. Genes under TATA box regulation are more able to respond rapidly (within a single generation) to environmental stress, show more variability in their expression range (phenotypic plasticity) compared to non-TATA regulated genes, and account for the appearance of stress induced phenotypes [22].

**Table 1.**
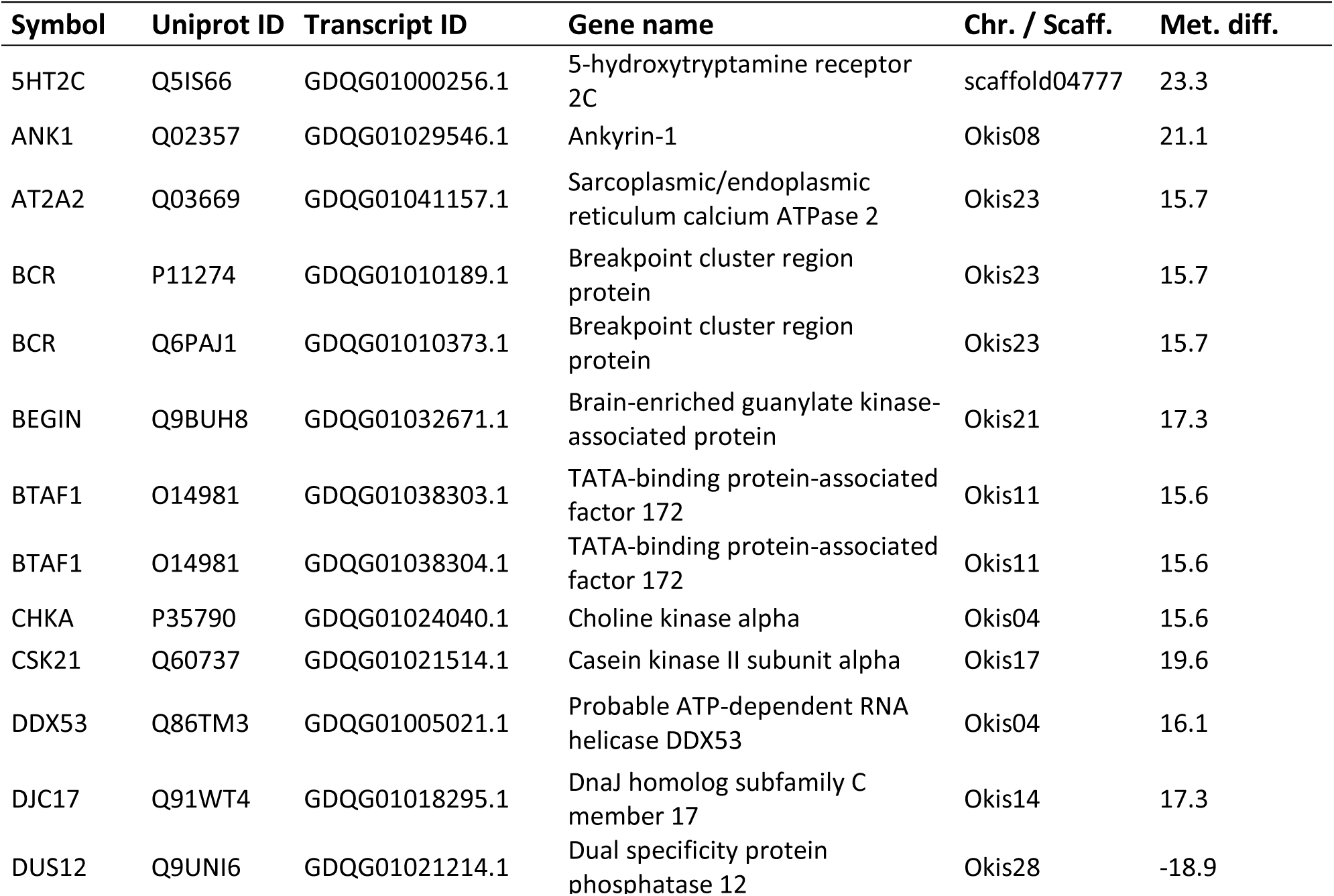
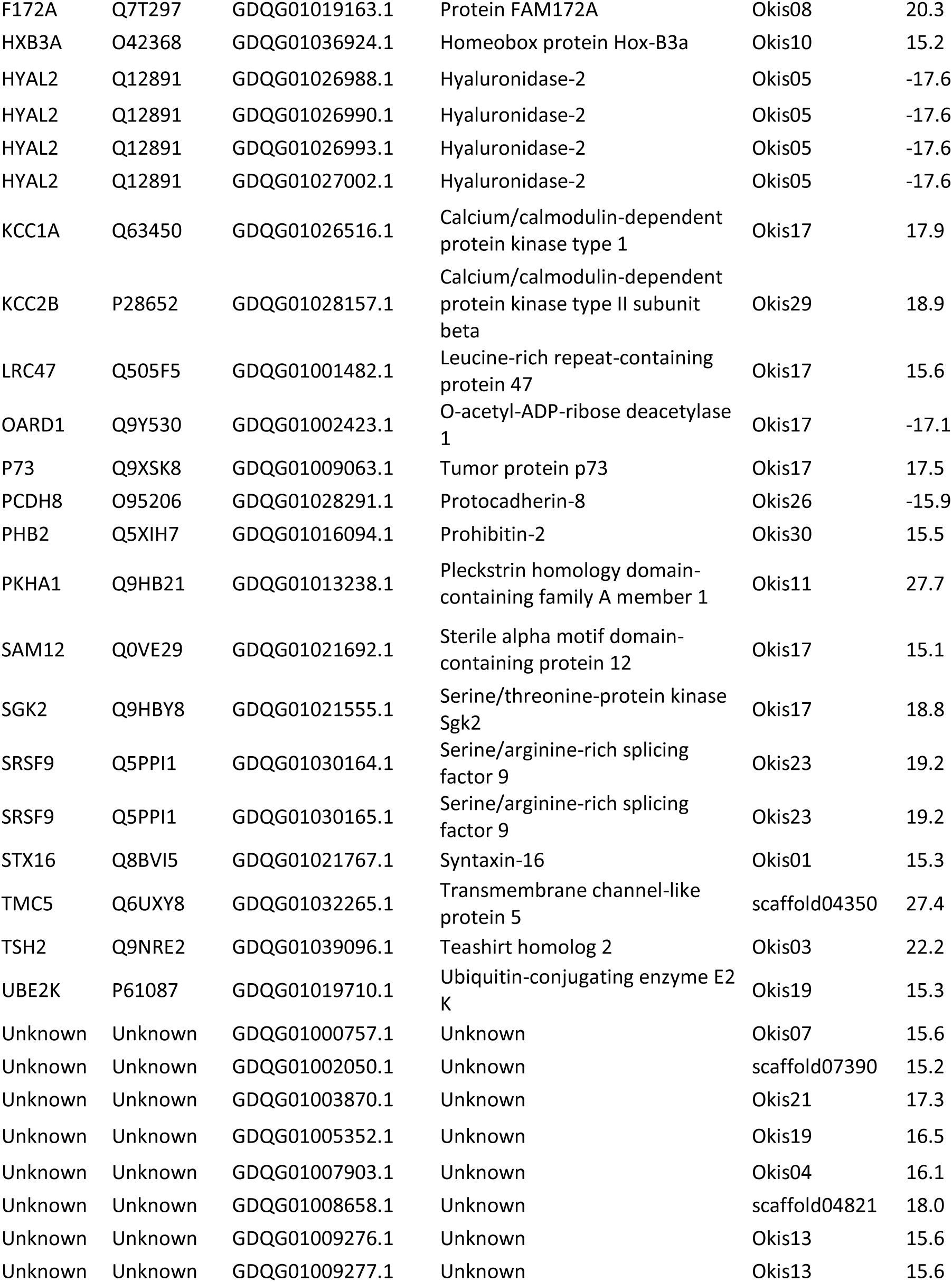
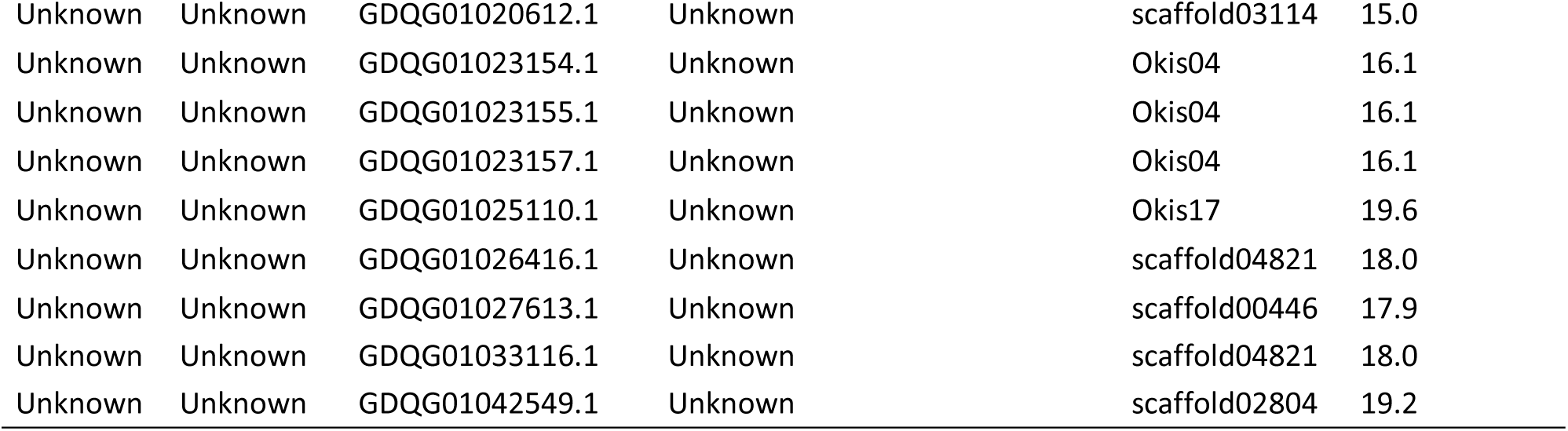
: Differentially methylated regions (DMRs) and their association with Uniprot entries between hatchery and wild smolt Coho Salmon. Annotation was based on a blastx approach against Uniprot-Swissprot database (e-value<10-6). Only significant DMRs were included (methylation difference between hatchery and wild (Meth. diff.) >15%; q-value< 0.01). Positive values are associated with hypermethylation relative to natural origin salmon. Transcript IDs correspond to the multi-tissue reference transcriptome for the Coho Salmon [14]. Each region represents a 1000 bp portion of one of the 30 chromosomes (Okis; Chr.) or additional scaffolds (scaffold; Scaff.) from the draft Coho Salmon genome assembly (GenBank assembly accession: GCA_002021735.1).

| Symbol | Uniprot ID | Transcript ID | Gene name | Chr. / Scaff. | Met. diff. |
| --- | --- | --- | --- | --- | --- |
| 5HT2C | Q5IS66 | GDQG01000256.1 | 5-hydroxytryptamine receptor 2C | scaffold04777 | 23.3 |
| ANK1 | Q02357 | GDQG01029546.1 | Ankyrin-1 | Okis08 | 21.1 |
| AT2A2 | Q03669 | GDQG01041157.1 | Sarcoplasmic/endoplasmic reticulum calcium ATPase 2 | Okis23 | 15.7 |
| BCR | P11274 | GDQG01010189.1 | Breakpoint cluster region protein | Okis23 | 15.7 |
| BCR | Q6PAJ1 | GDQG01010373.1 | Breakpoint cluster region protein | Okis23 | 15.7 |
| BEGIN | Q9BUH8 | GDQG01032671.1 | Brain-enriched guanylate kinase-associated protein | Okis21 | 17.3 |
| BTAF1 | O14981 | GDQG01038303.1 | TATA-binding protein-associated factor 172 | Okis11 | 15.6 |
| BTAF1 | O14981 | GDQG01038304.1 | TATA-binding protein-associated factor 172 | Okis11 | 15.6 |
| CHKA | P35790 | GDQG01024040.1 | Choline kinase alpha | Okis04 | 15.6 |
| CSK21 | Q60737 | GDQG01021514.1 | Casein kinase II subunit alpha | Okis17 | 19.6 |
| DDX53 | Q86TM3 | GDQG01005021.1 | Probable ATP-dependent RNA helicase DDX53 | Okis04 | 16.1 |
| DJC17 | Q91WT4 | GDQG01018295.1 | DnaJ homolog subfamily C member 17 | Okis14 | 17.3 |
| DUS12 | Q9UNI6 | GDQG01021214.1 | Dual specificity protein phosphatase 12 | Okis28 | -18.9 |
| F172A | Q7T297 | GDQG01019163.1 | Protein FAM172A | Okis08 | 20.3 |
| HXB3A | O42368 | GDQG01036924.1 | Homeobox protein Hox-B3a | Okis10 | 15.2 |
| HYAL2 | Q12891 | GDQG01026988.1 | Hyaluronidase-2 | Okis05 | -17.6 |
| HYAL2 | Q12891 | GDQG01026990.1 | Hyaluronidase-2 | Okis05 | -17.6 |
| HYAL2 | Q12891 | GDQG01026993.1 | Hyaluronidase-2 | Okis05 | -17.6 |
| HYAL2 | Q12891 | GDQG01027002.1 | Hyaluronidase-2 | Okis05 | -17.6 |
| KCC1A | Q63450 | GDQG01026516.1 | Calcium/calmodulin-dependent protein kinase type 1 | Okis17 | 17.9 |
| KCC2B | P28652 | GDQG01028157.1 | Calcium/calmodulin-dependent protein kinase type II subunit beta | Okis29 | 18.9 |
| LRC47 | Q505F5 | GDQG01001482.1 | Leucine-rich repeat-containing protein 47 | Okis17 | 15.6 |
| OARD1 | Q9Y530 | GDQG01002423.1 | O-acetyl-ADP-ribose deacetylase 1 | Okis17 | -17.1 |
| P73 | Q9XSK8 | GDQG01009063.1 | Tumor protein p73 | Okis17 | 17.5 |
| PCDH8 | O95206 | GDQG01028291.1 | Protocadherin-8 | Okis26 | -15.9 |
| PHB2 | Q5XIH7 | GDQG01016094.1 | Prohibitin-2 | Okis30 | 15.5 |
| PKHA1 | Q9HB21 | GDQG01013238.1 | Pleckstrin homology domain-containing family A member 1 | Okis11 | 27.7 |
| SAM12 | Q0VE29 | GDQG01021692.1 | Sterile alpha motif domain-containing protein 12 | Okis17 | 15.1 |
| SGK2 | Q9HBY8 | GDQG01021555.1 | Serine/threonine-protein kinase Sgk2 | Okis17 | 18.8 |
| SRSF9 | Q5PPI1 | GDQG01030164.1 | Serine/arginine-rich splicing factor 9 | Okis23 | 19.2 |
| SRSF9 | Q5PPI1 | GDQG01030165.1 | Serine/arginine-rich splicing factor 9 | Okis23 | 19.2 |
| STX16 | Q8BVI5 | GDQG01021767.1 | Syntaxin-16 | Okis01 | 15.3 |
| TMC5 | Q6UXY8 | GDQG01032265.1 | Transmembrane channel-like protein 5 | scaffold04350 | 27.4 |
| TSH2 | Q9NRE2 | GDQG01039096.1 | Teashirt homolog 2 | Okis03 | 22.2 |
| UBE2K | P61087 | GDQG01019710.1 | Ubiquitin-conjugating enzyme E2 K | Okis19 | 15.3 |
| Unknown | Unknown | GDQG01000757.1 | Unknown | Okis07 | 15.6 |
| Unknown | Unknown | GDQG01002050.1 | Unknown | scaffold07390 | 15.2 |
| Unknown | Unknown | GDQG01003870.1 | Unknown | Okis21 | 17.3 |
| Unknown | Unknown | GDQG01005352.1 | Unknown | Okis19 | 16.5 |
| Unknown | Unknown | GDQG01007903.1 | Unknown | Okis04 | 16.1 |
| Unknown | Unknown | GDQG01008658.1 | Unknown | scaffold04821 | 18.0 |
| Unknown | Unknown | GDQG01009276.1 | Unknown | Okis13 | 15.6 |
| Unknown | Unknown | GDQG01009277.1 | Unknown | Okis13 | 15.6 |
| Unknown | Unknown | GDQG01020612.1 | Unknown | scaffold03114 | 15.0 |
| Unknown | Unknown | GDQG01023154.1 | Unknown | Okis04 | 16.1 |
| Unknown | Unknown | GDQG01023155.1 | Unknown | Okis04 | 16.1 |
| Unknown | Unknown | GDQG01023157.1 | Unknown | Okis04 | 16.1 |
| Unknown | Unknown | GDQG01025110.1 | Unknown | Okis17 | 19.6 |
| Unknown | Unknown | GDQG01026416.1 | Unknown | scaffold04821 | 18.0 |
| Unknown | Unknown | GDQG01027613.1 | Unknown | scaffold00446 | 17.9 |
| Unknown | Unknown | GDQG01033116.1 | Unknown | scaffold04821 | 18.0 |
| Unknown | Unknown | GDQG01042549.1 | Unknown | scaffold02804 | 19.2 |

### No evidence for genome-wide differentiation between hatchery and natural origin salmon

We mapped the trimmed reads to the masked draft Coho Salmon genome assembly and identified 15,044 SNP markers (other than C-T polymorphism) meeting stringent filtering criteria and spread across the genome. The PCoA was produced on a Euclidean distance matrix of the 15,044 markers. Because no axis could be selected according to the broken-stick distribution, we selected all axes explaining at least 2.75% of the variation (10 axes explaining 33.9% of the variance), as previously performed with epigenetic markers [13]. A distance-based redundancy analysis (db-RDA) was produced on the genetic variation explained by these PCoA factors (response matrix) with river of origin, rearing environment and sex as explaining variables. The model was significant with an adjusted R^2^ of 0.18 (Figure 3). Both river of origin and sex were significant, whereas no significant effect was detected for rearing environment (Figure 3). No significant outlier with a genome-scan approach (Bayescan v2.0 [23]) was detected between sexes (Figure S2). Moreover, an AMOVA revealed no significant genome wide difference between rearing environments within river (Fst=0.005 and 0.002, for Capilano River and Quinsam Rivers, respectively; p-value>0.05) while the net difference between rivers was highly significant (mean Fst=0.038 ± 0.003; p-value<0.001; Table S2) [24]. Additionally, heterozygosity and inbreeding values (*G*IS) were not significantly different between rivers or between hatchery and natural origin fish (Table S3). Lastly, we used both a standard Bayescan genome-scan method for detecting outliers of large effect [23] and a Random Forest approach accounting for population structure, which allows detecting signals of polygenic selection. This statistical framework recently allowed detection of parallel polygenic selection between habitats within the panmictic North Atlantic eels (*Anguilla sp*.) and associated genetic variation with migration run timing in Chinook salmon (*Oncorhynchus tshawytscha*) populations [25–27]. No outlier (FDR>0.05) was detected between hatchery and natural origin fish using Bayescan v2.0 (Figure S3), whereas Random Forest identified 114 covarying markers, distributed over the 30 chromosomes. We used permutations (n = 1000, Supplementary experimental procedures) to assess whether a signal of apparent polygenic selection similar to the one that was detected could be obtained by chance (e.g. due to genetic drift or sampling error). Permutations reveal that similar pattern of apparent polygenic selection according to the distributions of the out-of-bag (OOB) errors could indeed be obtained by chance alone (Figure S4). Altogether population genomics analyses confirmed the prediction that hatchery and natural origin salmon belong to a single panmictic population within a given river. Our results cannot rule out that selection within one generation has caused changed in allele frequency between hatchery and natural origin fish in genome regions that were not screened. Nevertheless, they indicate that such an effect would be modest relative to parallel differences observed at the epigenetic level.

**Figure 3:**
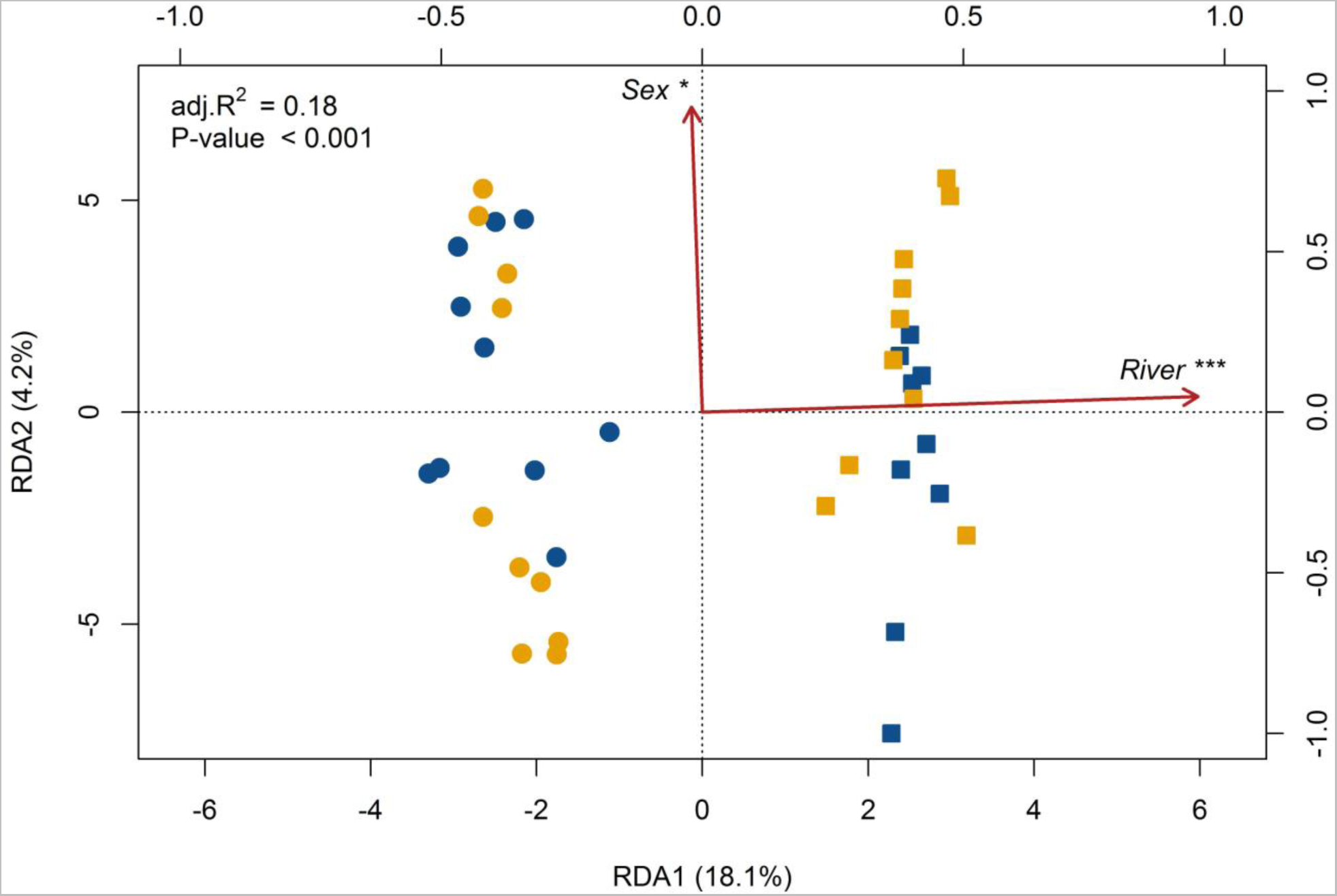
Distance-base redundancy analysis (db-RDA) performed on the total filtered 15,044 SNPs identified. Symbols represent rivers: Capilano (circle) and Quinsam (square). Colors represent captivity treatment: hatchery (blue) and wild (yellow). The db-RDA was globally significant and explained 18% of all SNPs variation (adj.R^2^=0.18). River of origin and sex explained significantly 16% and 2% of the variation after controlling for each other with subsequent partial db-RDAs. Asterisks represent p-value<0.001 (***) and p-value<0.05 (*) related to the explanatory factors.

## Discussion

The decline of many wild stocks of Pacific salmon encouraged the development of conservation hatcheries for enhancement. However, the hatchery environment during early life stages induces significant differences in the biology, physiology and behaviour, and ultimately in the fitness of hatchery-born fish [28]. Hatchery fish show higher reproductive success than their wild counterparts in hatchery conditions but lower success when released in the wild with an accumulative impact over a generation, advocating for inadvertent domestication effects occurring after a single generation of hatchery rearing [4,5,28]. Recent work provided evidence for a pronounced difference in gene expression between wild and hatchery fish after one year of captivity despite no significant differences at the genome level [3], as also reported between recently domesticated (five generations) Atlantic Salmon and their wild congeners [29]. However, epigenetic variation was recently associated with rapid adaptation to different natural environments (salt- vs fresh-water) in the Threespine Stickleback (*Gasterus aculeatus*) [30]. Here, for the first time, our results support the hypothesis that epigenetic modifications induced by hatchery rearing may represent a potential explanatory mechanism for rapid change in fitness related traits previously reported in salmonids. These similar epigenetic modifications were induced independently in two genetically distinct populations and in apparent absence of overall neutral and adaptive variation between hatchery and natural origin fish in these systems. This demonstrates that rapid epigenetic modifications are induced every generation during early development in the hatchery environment.

Indeed, combining a canonical multivariate approach (db-RDA) and pairwise Fst estimates, we found no significant evidence for genetic differentiation between hatchery and natural origin salmon, whereas genetic differentiation was highly significant between rivers systems. These results confirm that hatchery and natural origin fish belong to a single panmictic population, as predicted based on the hatchery programs applied in these rivers. These “integrated programs” are based on local populations and involve spawning in hatchery and natural environments. Hatchery and natural origin fish in each river are not kept separate, thus hatchery origin fish spawn in both the hatchery and the natural habitat as do natural origin fish, which can maintain high gene flow in the whole system.

Furthermore, no difference in genetic diversity (heterozygosity or inbreeding coefficient) was observed between hatchery and natural origin salmon, hence not supporting the hypothesis of increased probability of inbreeding depression in hatchery fish for the populations we studied. Finally, we found no evidence of either large effect or polygenic selection acting between hatchery and wild samples when using either a standard genome scan approach or statistical framework appropriate for investigating effect of weak selection in multiple regions of the genome. Therefore, our work corroborates recent findings on juvenile Steelhead Trout (*Oncorhynchus mykiss*) showing that only a single generation in captivity induced differences on the expression of hundreds of genes in offspring reared in identical environments, without a noticeable overall genetic difference (Fst=0.008; [3]).

In contrast to the apparent absence of significant genetic differences, our results revealed highly significant differences in methylation profiles between hatchery and natural origin salmon that were as pronounced as those observed between populations from different rivers. Our results differ from a study that compared hatchery-born and wild Steelhead Trout where no significant difference in methylation profiles was observed [12]. It may be that the impact of rearing environment on epigenetic modifications differs among species. It also may be that the negative result for trout was due to the lower resolution of the method available at that time, and indeed, the authors suggested that limited epigenetic differences between hatchery and wild fish could not be ruled out [12]. With a different approach offering substantial increase in genomic resolution, we found evidence for a highly significant effect of hatchery-rearing on DNA methylation profiles in many regions of the Coho Salmon epigenome, when controlling for population structure. Moreover, our results revealed that the same epigenetic modifications developed in parallel between the two independent study systems.

In Coho Salmon, it has been shown that hatchery fish are not as efficient as wild fish for rapid seawater acclimation[15]. In addition, acclimation to seawater induces profound, yet transient, changes in methylation levels in Brown Trout (*Salmo trutta* L.)[31]. We showed that genomic regions demonstrating differential methylation profiles between hatchery and wild salmon in both rivers were enriched for ion homeostasis and control of body fluid levels functions, adding growing evidence that hatchery rearing may affect the osmoregulatory process during smoltification. For instance, the serine/threonine-protein kinase (SGK2) is a potent stimulator of epithelial Na^+^ channels [32]. Similarly, seawater acclimation is associated with the level of SGK1 expression (no SGK2 or SGK3 ortholog present in the killifish genome) in the killfish (*Fundulus heteroclistus*) [33]. Considering the fundamental role of these biological functions during the smoltification (physiological adaptation to seawater), and migration of parr salmonids to the ocean [34], we propose that hatchery-induced epigenetic modifications during early developmental stages could be partly responsible for the saltwater acclimation deficiency reported in previous studies [35]. Moreover, neuromuscular communication, through regulation of Ca^2+^ levels, was among the biological functions showing the most pronounced differences between hatchery and natural origin salmon in both rivers. The enrichment was generally associated with a hypermethylation in hatchery fish, notably of a major regulator of motoneuron signal transmission through Ca^2+^levels (CAMK2). This observation strongly suggests an alteration of the neuromuscular communication that could reduce swimming performance as previously reported in hatchery-reared Coho Salmon [19]. Finally, although these results should be interpreted cautiously because they were limited to muscle tissue only, the enrichment for overall synaptic signal control functions raise the hypothesis that hatchery environment causes epigenetic modifications that may advocate a wealth of physiological and endocrinal differences. For instance, epigenetic differences we observed at some major neurological regulators such as HTR2C may play a role in the commonly observed behavioural differences between captive-reared and wild fish, such as increased aggressiveness, foraging, and boldness [30, 45–50]. This hypothesis could be tested by comparing methylation profiles in the brain between fish with different aggressiveness, foraging and boldness characteristics [41].

### Conclusions and implication for conservation and management

The reduced genome representation method used here and the fact that we could investigate only one tissue resulted in only a partial coverage of all possible epigenetic differences that may exist between hatchery and natural origin salmon. As such, our results should be interpreted as being conservative in reflecting the scale of epigenetic modifications incurred in the hatchery environment. Nevertheless, our results suggest that hatchery-rearing induces epigenetic variations that may alter the physiological (i.e. parr-to-smolt) transformation as well as the locomotor capacity that may result in reduced smolt fitness during juvenile seawater migration and ultimately, survival at sea. Whether or not the observed epigenetic modifications are inherited and be acted upon by natural selection cannot be answered from our results. Based on previous studies, it is reasonable to hypothesize that hatchery-induced epigenetic modifications during early developmental stages (post-fertilization and germ cell differentiation) almost certainly impact on lifelong phenotypic changes [42]. For conservation purposes, different practices in hatchery rearing are currently evaluated in order to circumvent the general observation that captive rearing reduces fitness in the wild. Alternative rearing practices may differ in environmental conditions (e.g. hatchery facilities or open lake), age at release (young fry or parr fish) or nutrition (supplemented or not by commercial food). For instance, previous work reported that salt-enriched food impact epigenetics and was correlated with a higher survival during sea transfer [31]. Clearly, improving our understanding of the dual role of genetic and epigenetic variation induced by captive rearing will contribute to development of the best practices for the management and conservation of salmonid fishes and possibly numerous other species that are managed through supplementation worldwide [43].

## Acknowledgements

We thank L. Benestan, B. Bougas, B. Boyle, A.-M. Dion Côté, C. Hernandez, M. Krick and E. Normandeau for laboratory and bioinformatics support. We thank O. Bichet, A.-L. Ferchaud, J.-S. Moore, C. Robert, Q. Rougemont, C. Rougeux and B. Sutherland for their comments. Computations were carried out on the supercomputer Colosse, Université Laval, managed by Calcul Québec and Compute Canada and on local servers (Katak). This research was carried out in conjunction with EPIC4 (Enhanced Production in Coho: Culture, Community, Catch), a project supported by the government of Canada through Genome Canada, Genome British Columbia, and Genome Quebec. The authors declare no conflicts of interest.

### Data accessibility

Sequences data are deposited on NCBI Short Read Archive database and will be made available upon acceptance (NCBI Bioproject: PRJNA389610).

### Authors contribution

J.LL, L.B designed the experiment. J.LL and M.L. analyzed the data and wrote the paper with L.B. J.LL performed the laboratory work. T.B, R.W, K.K designed and conducted the sampling. B.F.K, J.S.L and E.R provided the prepublication reference genome and re-sequencing data. All co-authors contributed substantially to revisions.

## Supporting Material

Appendix 1: Supplementary experimental procedures, figures S1-S4 and tables S1-S3

